# Multiparametric grading of glaucoma severity by histopathology can enable post-mortem substratification of disease state

**DOI:** 10.1101/2024.05.18.594740

**Authors:** Chuanxi Xiang, VijayKrishna Raghunathan, Yubin Qiu, Manisha Mehta, Cynthia L. Grosskreutz, Christopher W. Wilson, Ganesh Prasanna

## Abstract

Neurodegeneration in glaucoma patients is clinically identified through longitudinal assessment of structure-function changes, including intraocular pressure, cup-to-disc ratios from fundus images, and optical coherence tomography imaging of the retinal nerve fiber layer. Use of human post-mortem ocular tissue for basic research is rising in the glaucoma field, yet there are challenges in assessing disease stage and severity, since tissue donations with informed consent are often unaccompanied by detailed pre-mortem clinical information. Further, the interpretation of disease severity based solely on anatomical and morphological assessments by histology can be affected by differences in death-to-preservation time and tissue processing. These are difficult confounders that cannot be easily controlled. As pathogenesis and molecular mechanisms can vary depending on the stage and severity of glaucoma, there is a need for the field to maximize use of donated tissue to better understand the molecular mechanisms of glaucoma and develop new therapeutic hypotheses. Further, there is a lack of consensus around the molecular RNA and protein markers that can be used to classify glaucoma severity. Here, we describe a multiparametric grading system that combines structural measurements of the retinal nerve fiber layer with linear regression and principal component analyses of molecular markers of retinal ganglion cells and glia (RBPMS, NEFL, IBA1 and GFAP) to stratify post-mortem glaucoma eyes by the severity of disease. Our findings show that a quantitative grading approach can stratify post-mortem glaucoma samples with minimal clinical histories into at least three severity groups and suggest that this type of approach may be useful for researchers aiming to maximize insights derived from eye bank donor tissue.

## INTRODUCTION

Glaucoma is an optic neuropathy that affects approximately 80 million people worldwide, with up to 10% of glaucoma patients progressing to complete blindness [1–4]. It is characterized by progressive vision loss attributed to a loss of retinal ganglion cells (RGCs) and their axons, which form the optic nerve. Specifically, RGC loss accompanied by thinning of the retinal nerve fiber layer (RNFL) and cupping of the optic nerve head (ONH) are hallmarks of glaucomatous neuropathy [4, 5]. Several risk factors are associated with this disease; however, the only modifiable and treatable risk factor is elevated intraocular pressure (IOP). Current IOP-lowering treatments, either through pharmacological intervention (eye drops, intraocular drug implants, etc.) or surgical methods (e.g. trabeculectomy, goniotomy, etc.) have been shown to be effective in preventing vision loss [4]. However, many patients continue to experience vision loss despite well-controlled IOP clinically, suggesting that there are other processes in the inner retina and optic nerve head that could drive vision loss [6–8]. Some of these include neuroinflammation (astroglial and microglial activation, complement factors, etc.), vasoactive peptides and signaling molecules (e.g. endothelin-1 and nitric oxide interplay), and cytokines (e.g. TNFα) [9–11]. There is a need for novel therapeutics that can directly protect RGCs and axons to preserve vision in the long-term for glaucoma patients, in addition to IOP lowering therapies.

While glaucoma patients are diagnosed in the clinic using the structural characteristics mentioned above (including RNFL thinning and ONH cupping) by OCT, their clinical records are often either incomplete, sparse with long intervals between readings, variable, and/or non-transferrable between practices. This issue is exacerbated when patients sign up for willed-organ donation and donate their eyes post-mortem to qualified eye banks for research purposes, as records accompanying donor tissues may not include detailed deidentified clinical information. In addition, some post-mortem donor eyes (∼up to 20% in our experience) that are classified as non-glaucomatous according to medical records often show glaucoma-like characteristics after detailed histopathological examination, and thus may be mis-diagnosed. These confounders increase the challenge in accurately defining the glaucomatous disease stage of donated donor eye tissues after they have been processed for histopathology. Furthermore, pre-analytical variables such as fixatives used for histopathologic processing can introduce artifactual cupping of the optic disc. Appropriate staging and comparison to control samples is critical when evaluating changes in molecular pathways during the course of disease using techniques such as RNAScope or immunohistochemistry (IHC), as linking a particular pathway or potential therapeutic target to the stage of disease can be critical in determining when and where to therapeutically intervene. The use of post-mortem ocular tissue is critical in ophthalmic discovery research, as local assessment of potential drug targets at the site of disease pathogenesis is rarely possible in patients.

Recently, reports have described interesting observations regarding neuroinflammation in glaucomatous optic nerves/RGCs from human donor eyes and have compared and corroborated these findings to relevant preclinical models of glaucoma [12, 13]. These studies utilized histological assessments of ON atrophy, RGC numbers, and fibrosis in the ONH to define disease and to compare the extent of neuroinflammation, neutrophil infiltration and microglia/macrophage changes (CD163^+^ or CD68^+^ immunoreactivity), across glaucomatous vs. non-glaucomatous donor eyes. In the current study, we used hallmarks of RGC loss and gliosis to develop a quantitative multi-parametric and systematic grading system for post-mortem human glaucoma eye tissue sections. Structural measurements were made both in the central retina (macula) and temporal retina as well as the ONH, using the Bruch’s membrane opening (BMO) as the landmark. We then used linear regression and principal component analyses to study correlations between the RGC soma marker RBPMS [14], the RGC axonal marker neurofilament light chain (NEFL) [15, 16], the microglia/macrophage marker IBA1 [17], and the astrocyte marker glial fibrillary acidic protein (GFAP) [18, 19]. Our findings reveal that a quantitative structural and molecular marker grading approach can stratify post-mortem glaucoma samples into at least three severity groups, and this type of approach may be useful for researchers aiming to maximize insights derived from eye bank donor tissue.

## MATERIALS AND METHODS

### Human donor eye acquisition, processing, and IHC

Sixteen human eye globes (HEGs) with no history of glaucoma and twenty-two HEGs with an accompanying medical history of glaucoma but no detailed clinical information were collected from the Lion’s Eye Institute for Transplant & Research (Tampa, FL) between 2017 and 2022. Donor demographics and causes of death are listed in **Table 1**. The HEGs were enucleated within 6 hours post-mortem and evaluated as not meeting the criteria for transplantation. All samples were shipped in Modified Davidson’s Fixative (MDF) and fixed for 48 hours, and then transferred to 70% alcohol for 48-72 hours. HEGs were cut transversely in the pupillary-optic axis into 3 parts with the optic nerve and macula contained in the central part and then embedded in paraffin in a Tissue-Tek ® VIP™ processor and Tissue-Tek Tec (Sakura Finetek USA). 5-micron transverse sections with macula and the optic nerve were cut from the central part. These sections were then processed using standard dehydration, deparaffinization and rehydration procedures in and stained with hematoxylin and eosin (H&E). IHC was performed with antibodies (**Table 2**) against RBPMS, NEFL (RGC marker), GFAP (astrocyte marker) and IBA1 (microglia/macrophage marker) using a LeicaBond RX stainer (Leica Biosystems, Deer Park, IL, USA). Heat induced epitope retrieval was performed in a solution of EDTA based epitope retrieval solution (HIER, pH 9.0) (AR9640, LeicaBond) at 95°C for 20 minutes.

### Image analysis and quantification

IHC stained slides were scanned using an Aperio AT2 scanner (Leica Biosystems) and subsequently imaged and quantified using the HALO image analysis platform (Indica Labs, New Mexico USA). Bruch’s membrane opening (BMO) was used as a landmark for the optic nerve. IHC signals in the ganglion cell complex (GCC, including nerve fiber layer, ganglion cell layer, and inner plexiform layer) were collected from the central region (1 mm range from temporal BMO), peripheral retina (2 mm area beginning 7 mm from temporal BMO) and ONH (1 mm area below the line of the two sides of the BMO). RGC cells were counted from 5 mm range of the periphery (5 mm area beginning at 7 mm from temporal BMO) and 1 mm in the central retina (1mm from the temporal BMO) (**Supplemental Fig. 1**).

### Statistical analysis and glaucoma disease stratification

#### Descriptive statistics and hypothesis testing

Tissue regions positively stained for NEFL, GFAP, and IBA1 were expressed as a percentage of positively stained area per total area measured. Retinal ganglion cells are reported as number of RBPMS positive cells per 6 mm of length of the retina. GCC thickness was measured in the central temporal region and reported as µm. Differences in NEFL, GFAP, IBA1, RBPMS and retinal thickness are reported as scatter plots with individual values from each donor, with horizontal lines indicating the mean ± standard deviation. Comparison between glaucomatous and non-glaucomatous groups were performed by unpaired t-test with p<0.05 as being statistically significant (*p<0.05, **p<0.01, ***p<0.001, ****p<0.0001). Subsequently, due to the binary nature of the groups and in order to determine if the differences between the two groups were true, we performed the receiver operating characteristic (ROC) curve analysis. The degree of separability between sensitivity (true positive) and specificity (false positive) are indicated by the area under the curve (AUC) on the graphs. Next a linear regression analysis was performed to study the correlation between RBPMS:IBA1, RBPS:NEFL, RBPMS:GFAP, and IBA1:GFAP comparing glaucomatous and non-glaucomatous donors. Goodness of fit and correlation is expressed by the r^2^ value.

#### Disease stratification

In order to further understand the data, we sought to determine if performing multivariate analysis and data clustering could classify the data by disease severity. First, we performed principal component analysis (PCA) from a data matrix consisting of 38 samples (16 non-glaucoma, 22 glaucoma donors) and 4 variables (NEFL, RBPMS, GFAP, IBA1). The data matrix was centered and normalized to get a mean value of zero and a standard deviation of 1 for each variable. A correlation matrix was calculated, and the eigenvectors and corresponding eigenvalues were extracted. The relative percentage of variance of each eigenvalue was quantified and diagrammed in a Scree plot (**Fig. 8A**) [20]. The selection of the number of principal components used was performed using the Kaise-Guttman criteria[21], using the eigenvectors whose corresponding eigenvalues had a value greater than 1. Subsequently, the score matrix was calculated by multiplying these eigenvectors by the square root of the corresponding eigenvalue. Following PCA, the score matrix was rotated orthogonally using Varimax [22] to extract the latent factors. The factor loading matrix was obtained from the rotated factor score matrix by matrix operations. The factors were interpreted by associating the variables with highest weights in each factor and expressed as a score plot (**Fig. 8B**). PCA combined with factor analysis revealed that 3 predominant variables contributed to maximal variance in the samples (i.e. NEFL, RBPMS, and GFAP). Finally, by applying the Calinski-Harabasz criterion [23] for unbiased k-means clustering, we determined the optimal number of clusters for both non-glaucomatous and glaucomatous donors, using the 3 variables identified, and the data was clustered and represented as 3D plot.

## RESULTS

In order to determine the extent of glaucomatous damage in the posterior segment of donor eyes, we systematically evaluated histological features from 22 glaucomatous eyes (65-97 years old) and 16 age-matched non-glaucomatous control eyes. The histologic parameters evaluated were the thickness of the GCC, cupping of the optic disc, number of remaining RGC soma, immunoreactivity of NEFL in retinal and optic nerve sections, and glial activation (immunoreactivity for GFAP and IBA1). At this stage, since limited clinical and medical histories of the donors were available to us, the donors remained unstratified.

### Gross anatomical changes in retina, macula, and optic nerve head in glaucoma

Macular damage in samples with a positive medical history of glaucoma was observed by a discernible reduction in RGC cell numbers and density within the temporal macular region from H&E in 12 of 22 donors (**Fig. 1A**). In some donors, the RGC layer was completely lost in both temporal and nasal macular regions (**Supplemental Fig. 2A**) accompanied by local RGC loss in the central macula for one donor (**Supplemental Fig. 2B**). Cupping of the optic disc was observed in 11 of 22 glaucomatous donor eyes (**Fig 1B**). Remodeling of the lamina cribrosa (compression and disorganization of the prelaminar, lamina and post-lamina regions) was observed in 13 of 22 donor eyes. The extent of cupping and lamina cribrosa remodeling varied between donors and did not always correlate with RGC loss [24–27]. Given that sample fixation and processing for histology could confound accurate assessment of cupping, we did not consider cupping for additional sample classification or stratification.

**Figure 1:**
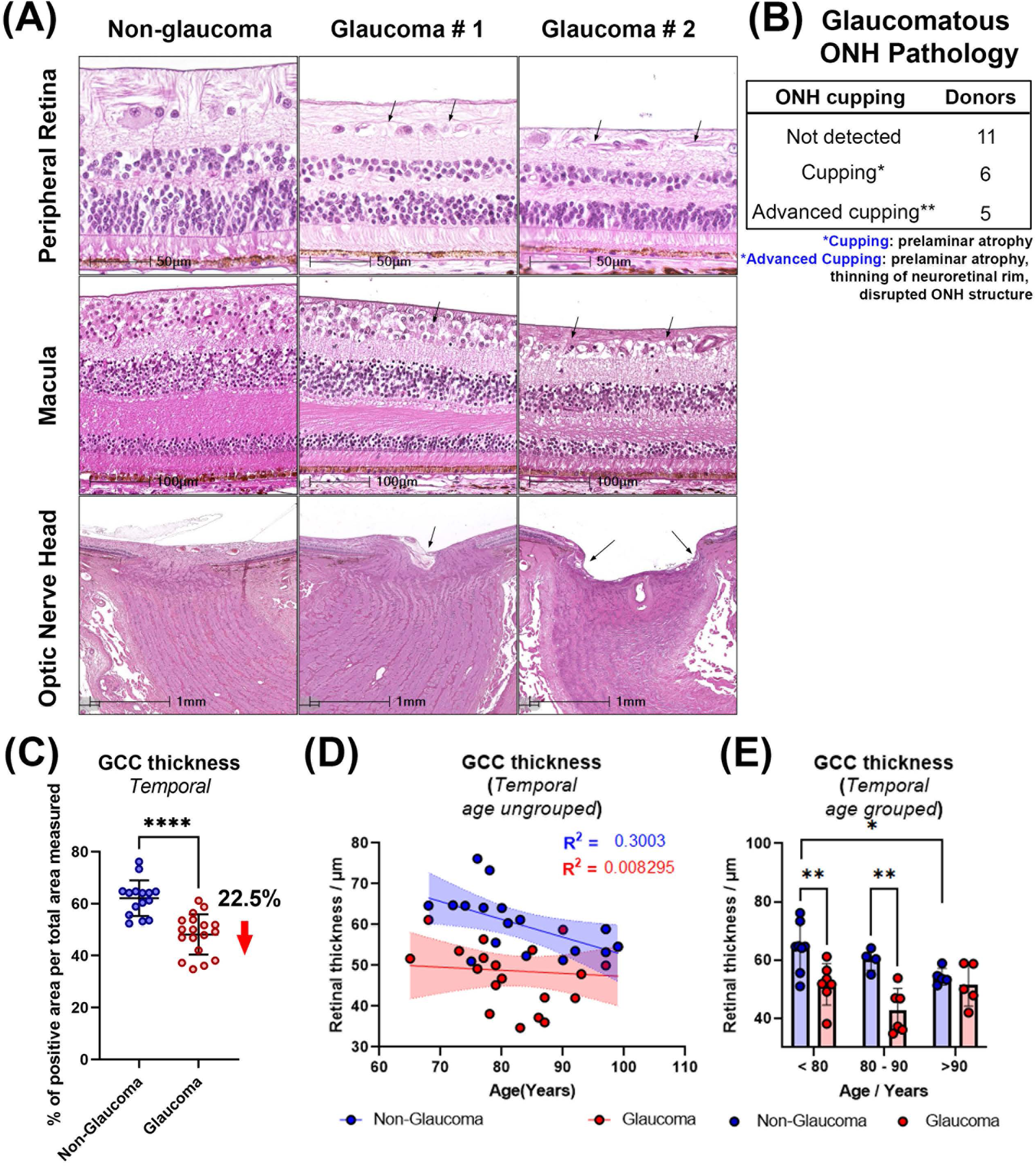
Pathological changes of human glaucomatous retina and ONH. **(A)** Histological changes demonstrating decrease in peripheral retinal thickness, macular RGC loss, reduced RGC cell density, GCC thickness, ONH cupping in glaucoma. Arrows in top and middle panels indicate thinning of the RNFL and arrows in the bottom panels demonstrate cupping. (**B**) ONH cupping or advanced cupping was often seen in glaucoma, which showed prelaminar atrophy, disrupted structures in optic nerve head and in optic nerve, and lamina cribrosa bowed posteriorly. **(C)** Scatter dot plot represents the individual values of temporal GCC thickness for non-glaucomatous and glaucomatous donor tissues. GCC thickness was significantly decreased by 22.5% in temporal peripheral retina of glaucomatous donors. Horizontal bars indicate mean ± standard deviation. ****p<0.0001, unpaired t-test. **(D)** GCC thickness decreases with age in non-glaucomatous donors (R^2^=0.3), but such a trend is not apparent in glaucomatous donors (R^2^=0.0). **(E)** Significant differences in GCC thickness are seen comparing glaucoma and non-glaucoma donors in patients who are below 80 years and between 80-90 years. No significant difference in GCC thickness is seen in patients > 90 years of age comparing glaucoma and non-glaucoma donors.

Next, we compared GCC thickness of non-glaucomatous and glaucomatous retinas from donors (**Fig. 1C-E**). Qualitative thinning of the peripheral retina (RGC loss, RFNL thinning) was observed by H&E staining (**Fig. 1A**) in 12 of 22 glaucomatous retinas. We next directly evaluated RGC soma loss in the peripheral retina comparing temporal to macular regions, and the nasal to central area (between macula and ONH). RGC loss was noted in both regions around the macula (the temporal and nasal) from 8 of 12 donors with retinal thinning, but only in the temporal retina of the remaining 4 donors with thinning. To confirm reduction of RNFL thickness in glaucoma, GCC thickness was measured in the temporal peripheral retina of all donor retinas (temporal to macula, 7 mm from BMO, **Supplemental Fig 1**). On average, the GCC thickness of non-glaucomatous eyes was 61.9 ± 1.7 µm (mean ± SEM for all numerical values thereafter unless stated otherwise) while it was significantly lower by 22.5% (49.2 ± 1.80 µm) in glaucomatous eyes (**Fig. 1C**). We observed that, in non-glaucomatous donors, GCC thickness trended towards a decrease with age (R^2^ = 0.3 **Fig. 1D**). This trend was absent in glaucomatous donors (R^2^ = 0.008), where GCC layer thinning had no relationship with age. When samples were stratified into 3 age groups (<80y, 80-90y, >90y) we observed a statistically significant reduction in GCC thickness when comparing eyes from non-glaucomatous donors over 90 years of age to those lesser than 80 years of age (Fig. 1E). There was no association with age in glaucomatous donors (**Fig. 1E**). Also, when compared with non-glaucomatous donors, GCC thickness was significantly lower in glaucomatous donors in both the <80y and 80-90y groups. No significant difference in GCC thickness was observed between the two groups in the >90y age group. Together, the data demonstrate that in this cohort of samples, disease state contributes to a greater thinning of the GCC in donors <90y of age.

### Loss of retinal ganglion cells in glaucoma

To confirm RGC loss observed by H&E staining, we immunolabelled tissues with the RGC specific marker, RBPMS. Similar to our prior assessments, we observed 5-7 layers of RBPMS+ RGCs in non-glaucomatous donors. However, in glaucomatous donors, most sections exhibited 2-5 RBPMS+ cell layers. In some cases, there were only 0-2 layers suggesting advanced severity (**Fig. 2A**). Further quantitative analysis was performed in the temporal peripheral retina (5 mm from the end of the macular outline) and nasal retina (1 mm BMO and macula) (**Supplemental Fig. 1**). We observed a 53.8% decrease in RGC density in the temporal peripheral retina from glaucomatous donors compared with non-glaucomatous donors (**Fig. 2B**).

**Figure 2:**
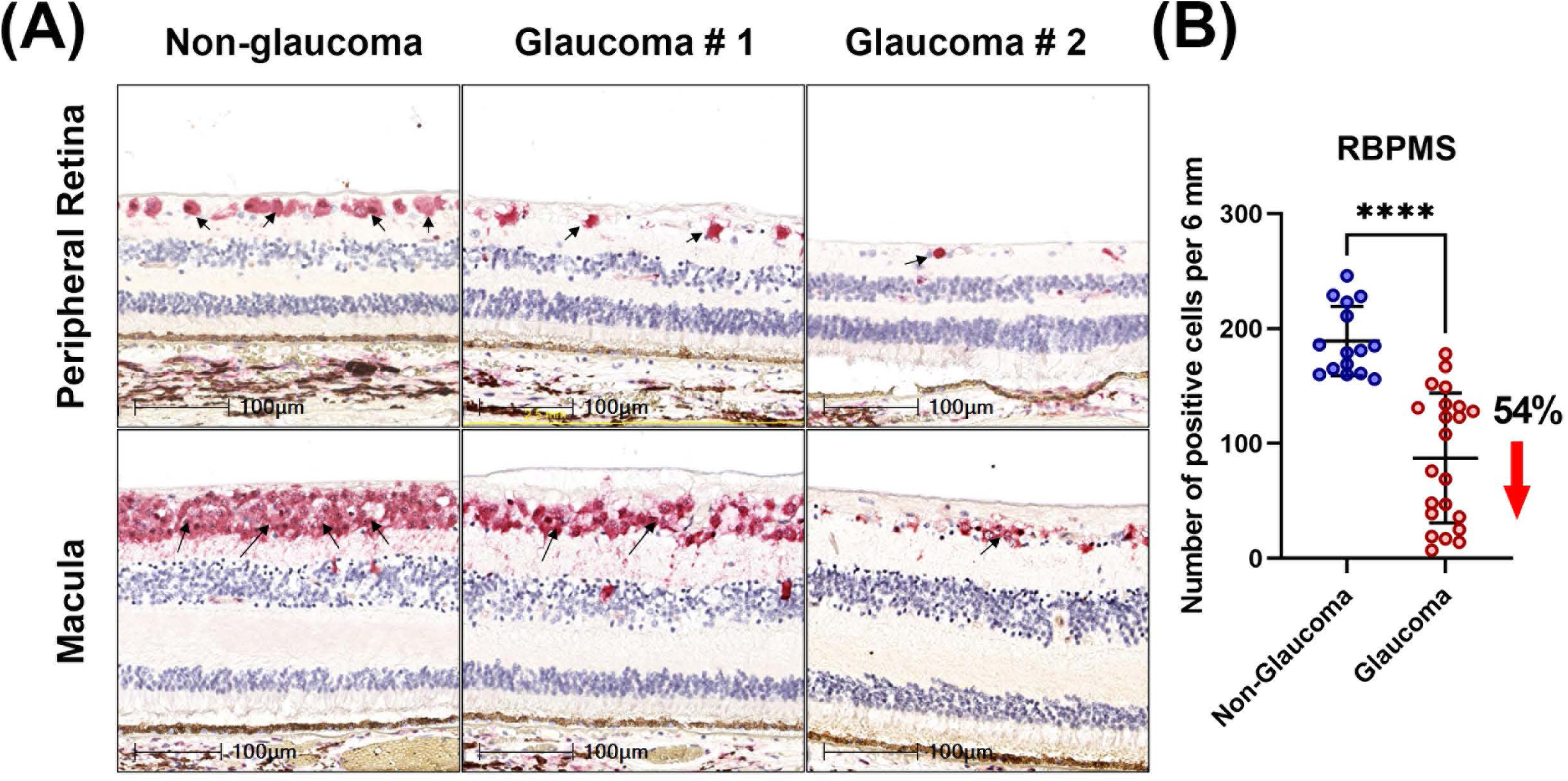
RGC loss in glaucomatous donors observed by loss of RBPMS positive cells. **(A)** Loss of RGCs in peripheral retina and macula was observed in glaucomatous donors; arrows indicate RBPMS positive cells. **(B)** Scatter dot plot represents the individual values of RBPMS-positive RGCs (RBPMS+ RGCs) for non-glaucomatous and glaucomatous donor tissues. Quantitative analysis demonstrated a 53.84% reduction in RBPMS+ RGCs in glaucomatous temporal peripheral retina, Horizontal bars indicate mean ± standard deviation. ****p<0.0001, unpaired t-test.

### Axonal and dendritic loss in glaucoma

Loss of axonal integrity and dendrites in the neural retina was determined by measuring the immunoreactivity of neurofilament light chain protein (NEFL) in both the peripheral temporal retina and ONH. Qualitative evaluation revealed a reduction in NEFL+ processes in the peripheral temporal neural retina accompanied by irregular distribution of NEFL in the glaucomatous ONH (**Fig. 3A**). Expression levels of NEFL were quantified within a 2 mm peripheral temporal region and 1 mm central region between the BMO and macula by normalizing the percentage positive immunolabel to total area measured. Normalized NEFL immunoreactivity in non-glaucomatous donor retina was 63.5 ± 2.9%, while it was significantly reduced to 36.2 ± 4.2% in glaucoma (**Fig. 3B**) demonstrating a net reduction of NEFL+ axons by 40%. The reduction in NEFL immunoreactivity in glaucomatous ONH was even greater (82.6%) (**Fig. 3B)**.

**Figure 3:**
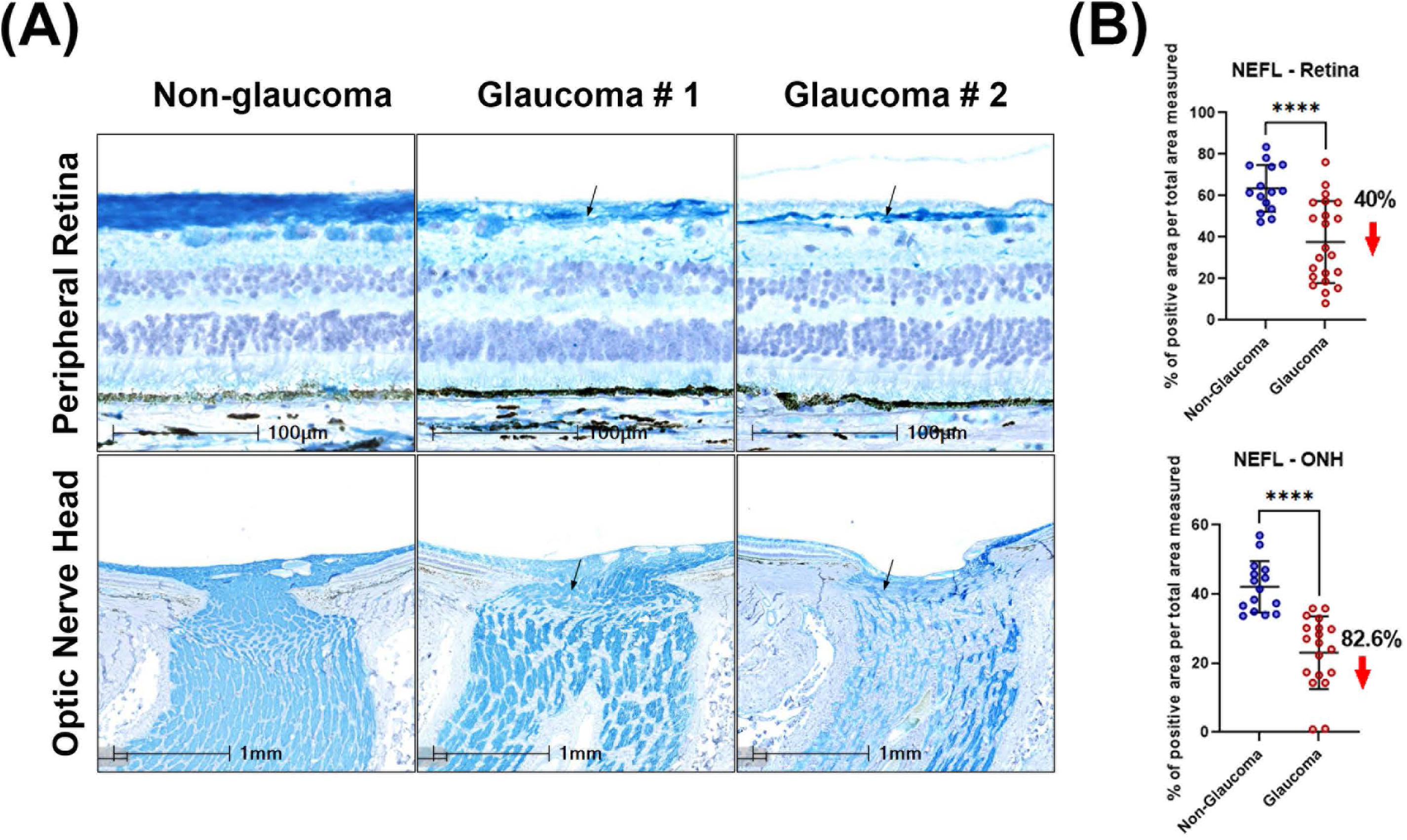
Axonal and dendritic loss of RGCs visualized by NEFL immunolabelling in the peripheral retina and ONH. **(A)** Decreased NEFL expression in temporal peripheral retina and ONH, and disorganization of NEFL immune staining was observed in glaucomatous tissues. Arrows indicate representative areas with loss of NEFL. **(B)** Scatter dot plot represents the individual values of neurofilament light chain (NEFL) for non-glaucomatous and glaucomatous donor tissues. Quantitation of NEFL staining showed a statistically significant reduction in the temporal peripheral retina (40%) and optic nerve head (82.6%) of glaucoma donors. Horizontal bars indicate mean ± standard deviation. ****p<0.0001, unpaired t-test.

### Glial activation in glaucoma

Qualitative assessment revealed increased immunoreactivity of GFAP in the glaucomatous peripheral retina accompanied by disorganized GFAP+ regions within the ONH, compared with non-glaucomatous donors. Specifically, in non-glaucomatous tissues, when GFAP+ signals were observed, these were localized to inner layers of the retina (especially in RNFL and RGCL) and around blood vessels. However, in glaucomatous tissues, the intensity of GFAP labelling was significantly higher and extended from the inner retina to the inner nuclear layer (INL), outer plexiform layer (OPL), and in some cases, to the outer nuclear layer (ONL) (**Fig. 4A**). Also, increased GFAP intensity was observed in glaucomatous ONH throughout the peripapillary, prelaminar and lamina cribrosa regions. Disorganized GFAP staining patterns were also observed in post-laminar regions of glaucomatous ONH (**Fig. 4A**). Using a similar quantitative approach to NEFL measurement, we report a 73.3% increase in normalized GFAP-immunoreactivity in glaucomatous temporal peripheral retina compared with non-glaucomatous donor retinas (**Fig. 4B**). Similar increases in GFAP immunoreactivity was seen in the ONH (75.8%).

**Figure 4:**
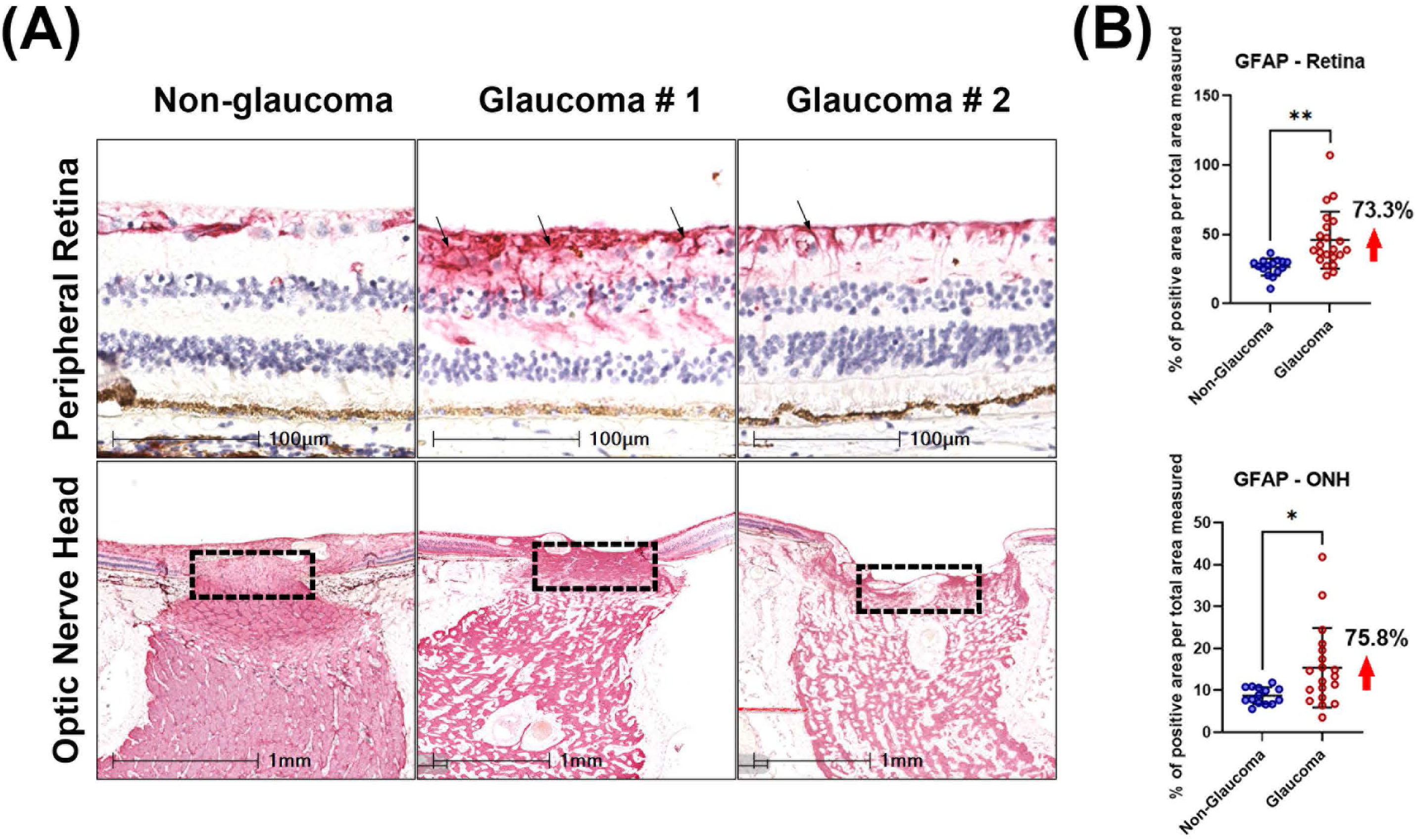
Detection of activated astrocytes by GFAP immunoreactivity. **(A)** Compared with non-glaucomatous donors, upregulation of GFAP was detected in glaucomatous peripheral retina and ONH. Arrows indicate representative GFAP immunoreactivity. The dotted boxes on the bottom panel highlight the region of the ONH quantified in Figure 4B. **(B)** Scatter dot plot represents the individual values of reactive astrocytes (GFAP+ cells) for non-glaucomatous and glaucomatous donor tissues assessed by histology. Quantitative analysis showed that GFAP was significantly upregulated in the temporal peripheral retina (73.3%) and optic nerve head (75.8%) of glaucoma donors. Horizontal bars indicate mean ± standard deviation. **p<0.01, unpaired t-test.

Next, we evaluated the number of microglia present in the tissue by immunolabelling for IBA1+ cells. In glaucomatous retinas, IBA1+ microglia were mostly present in the RNFL, ganglion layer, IPL and INL (**Fig. 5A**). Dual staining of IBA1 and RBPMS detected IBA1+ cells to be closely associated with RBPMS+ RGCs (**Supplemental. Fig 3**). A significantly greater cell density (% positive cells per total area measured) of IBA1+ microglia was observed in glaucomatous retinas (7.8 ± 3.6 %) compared with non-glaucomatous tissues (3.7 ± 1.1%; **Fig. 5B**) reflecting a net increase by a factor of 100%. In the optic nerve head (within the first 1 mm below B, a significant increase in IBA1 positive cell density was observed in glaucomatous tissues compared with non-glaucomatous tissues (a factor of 70.9%; **Fig. 5B**).

**Figure 5:**
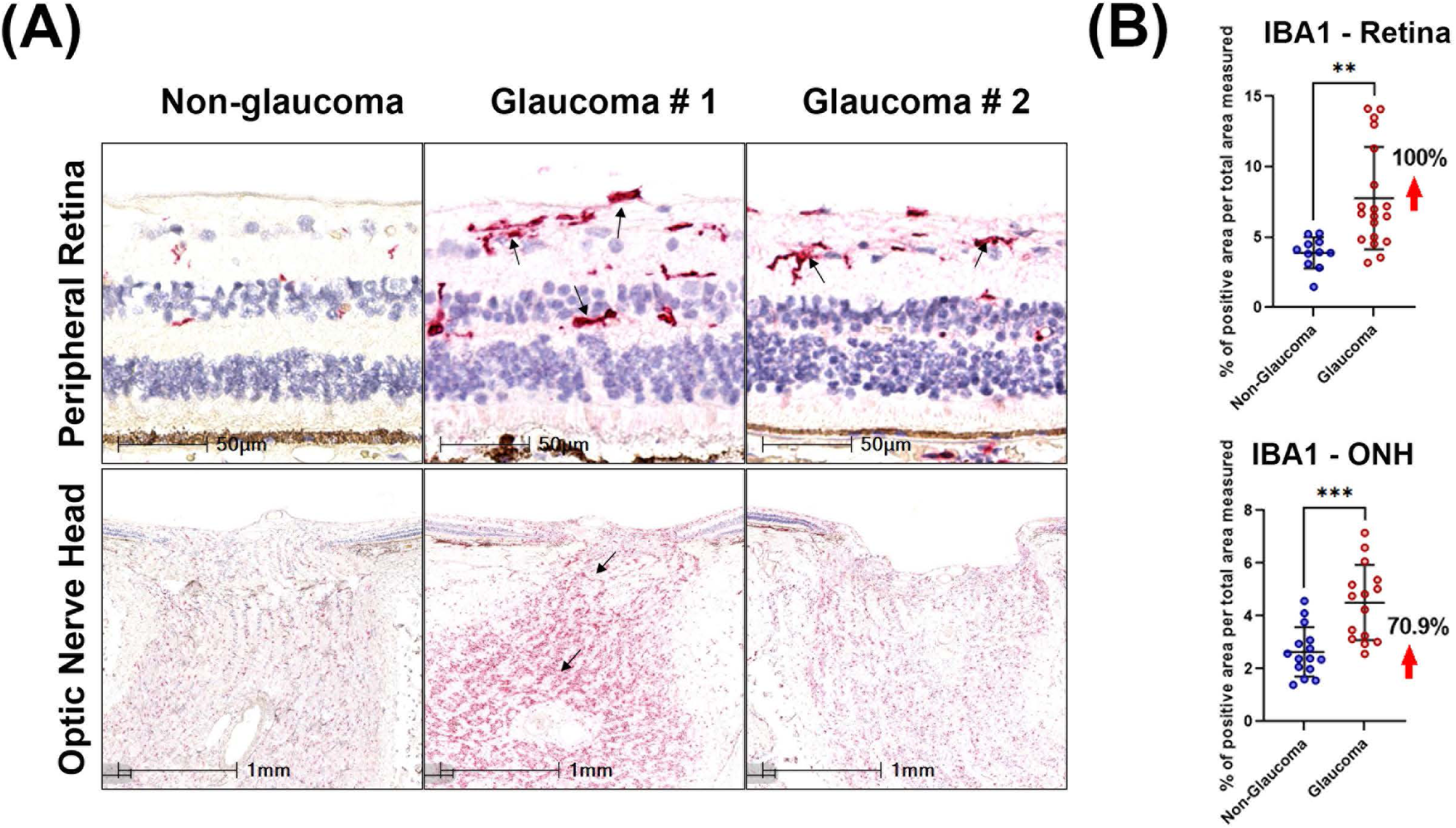
Detection of microglia by IBA1 immunoreactivity. **(A)** IBA1 was upregulated in glaucomatous peripheral retina and ONH compared with non-glaucomatous donors. Arrows indicate IBA1 positive cells in the retina and increased IBA1 immunoreactivity in the ONH. **(B)** Scatter dot plot represents the individual values of IBA1+ cells for non-glaucomatous and glaucomatous donor tissues assessed by histology. Quantitative analysis of IBA1 positivity showed that IBA1 was statistically upregulated in the temporal peripheral retina (100%) and optic nerve head of (70.9%) glaucoma samples. Horizontal bars indicate mean ± standard deviation. **p<0.01, unpaired t-test.

### Stratification of disease severity by multivariate analysis

After assessing individual markers, we sought to determine if there was any correlation between them. First, we asked if there was any correlation between protein marker staining between the retina and optic nerve head tissues. This analysis was performed only on sections from donors where both the retina and optic nerve were present. Spearman’s analysis demonstrated a statistically significant correlation for GFAP and NEFL immunolabelling between the retina and optic nerve head, but no statistical significance was observed for IBA1 labelling between the two tissues (**Fig. 6**). For all subsequent analyses, measurements from only the retinal tissue were used.

**Figure 6:**
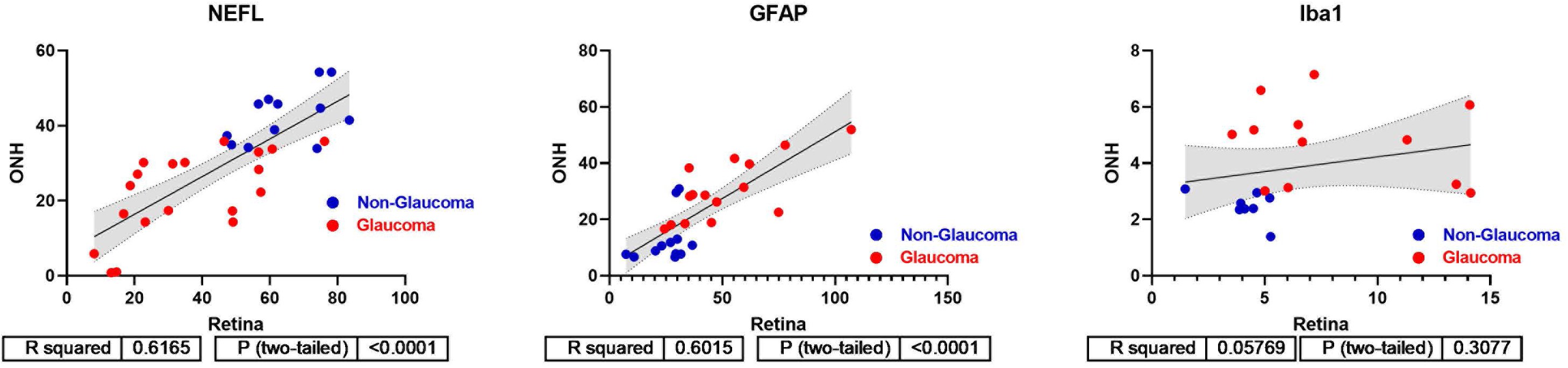
Correlation analysis between retina and optic nerve head. Spearman’s correlation and linear regression was performed for NEFL, GFAP and Iba1 expression comparing retina and ONH measurements for glaucoma and non-glaucoma donors. NEFL and GFAP exhibit strong correlation between retina and ONH measures, but this is not the case for IBA1.

Since loss of RGCs is considered a hallmark of disease, we performed all our Spearman correlation analyses against quantitative RBPMS+ immunoreactivity in the temporal retina. We observed a positive correlation between loss of NEFL+ immunoreactivity and loss of RBPMS+ RGC cell density (R^2^=0.8097; Spearman correlation, **Fig 7A**) in glaucomatous tissues. This was a significant increase when comparing the same parameters in non-glaucomatous donors (R^2^=0.3118; Spearman correlation). No significant correlation was observed comparing RBPMS and GFAP or IBA1 levels in either glaucoma or non-glaucomatous donors. In the subset of donors where IBA1 positive immunoreactivity was observed, no relationship between GFAP and IBA1 immunostaining was observed in either glaucomatous or non-glaucomatous donors (**Fig. 7B**); although the variability in staining for both parameters was significantly greater for glaucomatous donors than non-glaucomatous donors.

**Figure 7:**
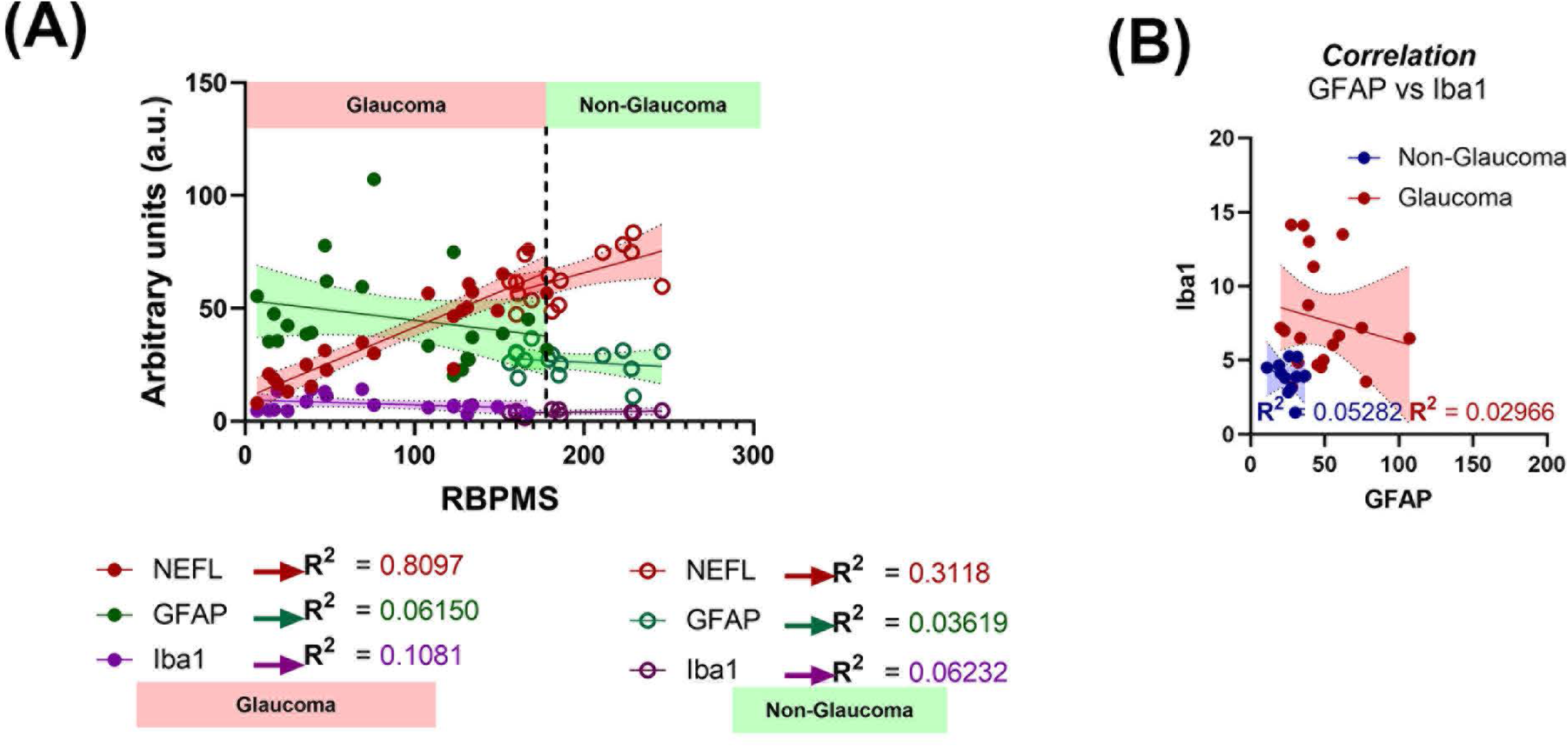
Spearman’s correlation analysis. **(A)** Spearman’s correlation demonstrated maximal correlation between RBPMS and NEFL in glaucomatous donors, with minimal correlation seen when comparing other markers in glaucomatous and non-glaucomatous donor tissues. **(B)** No statistically significant correlation was observed when comparing GFAP and IBA1 immunoreactivity in the retina for both non-glaucomatous and glaucomatous donors.

**Figure 8:**
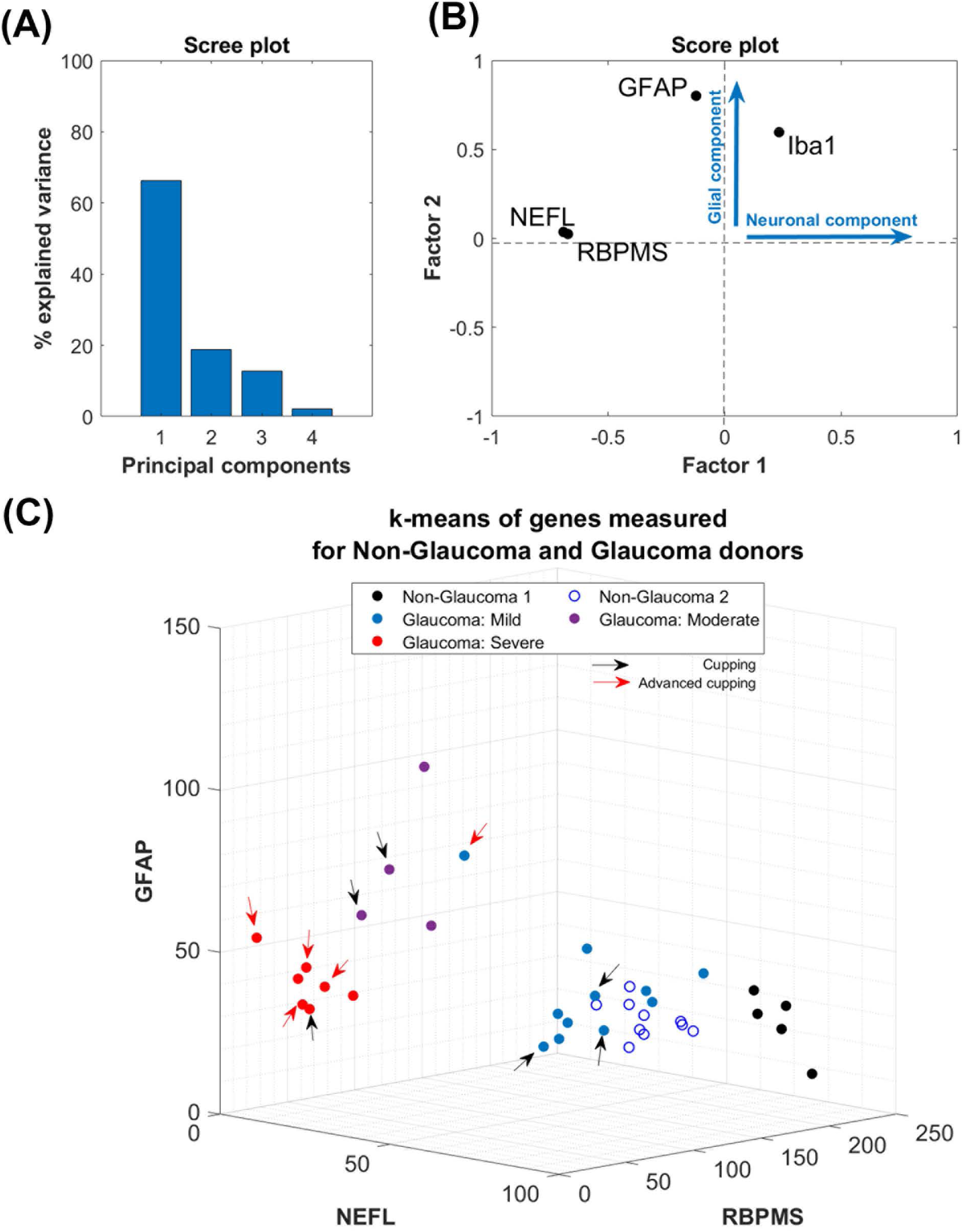
Multivariate statistical analysis. **(A)** Principal component analysis for GFAP, RBPMS, NEFL and IBA1 immunoreactivity demonstrated that > 85% of the variance in the data was contributed by the first 2 principal components, as indicated in the Scree plot. **(B)** Score plot from factor analysis showing that maximal variance was weighted by 3 factors: RBPMS, NEFL and IBA1, with the former 2 representing neuronal components, and the latter representing a glial component. **(C)** Cluster analysis using the principal components. Severity of glaucoma was assessed by applying the Calinski-Harabasz criterion for unbiased k-means clustering using the expression values for RBPMS, NEFL, and GFAP positive staining. Non-glaucomatous donors clustered in 2 distinct groups while glaucomatous tissues clustered in 3 distinct groups (mild, moderate, and severe). Arrows as indicated in the figure identify tissues with qualitative cupping of the ONH observed through histology.

Next, to determine the extent that donor variability contributed to the significance in quantitative measurements, and to compare sensitivity vs. specificity of the outcomes comparing glaucomatous and non-glaucomatous donors, we performed receiver operating characteristic (ROC) curve analysis (**Supplemental Fig. 4**). ROC analysis revealed that the area under the curve was the greatest for RBPMS (AUC = 0.9667) and those for NEFL (AUC = 0.8485), GFAP (AUC = 0.8697), and IBA1 (AUC = 0.8804) were significant reinforcing the results that the quantitative changes observed in glaucoma were true positives.

Since variability was observed both from the scatter and ROC plots, we hypothesized that these changes in temporal retinal NEFL (axonal & dendritic loss), RBPMS (RGC loss), and GFAP (glial activation) levels may be reflective of disease stage. PCA and factor analysis of the data revealed that the first two principal components with 3 variables (NEFL, RBPMS, and GFAP) contributed to the maximal variance (>85%) in the data (**Fig. 8A, B**). Using the quantitative measurements for these 3 variables, unbiased k-means clustering revealed 3 clusters for glaucoma samples and 2 clusters for non-glaucoma donors (**Fig. 8C**).

The two non-glaucoma clusters were predominantly factored on possessing a high level of RBPMS+ and NEFL+ staining, and a low proportion of GFAP+ glial cells. For glaucomatous tissues, the clustering was parameter dependent. Donor sections with the highest level of RBPMS+ NEFL+ staining and least glial proliferation/GFAP positivity were categorized as mild glaucoma. Donor samples with the greatest increase in GFAP+ cells but moderate loss of NEFL+ RBPMS+ staining were classified as moderate glaucoma. Finally, donors with greatest loss of RBPMS+ NEFL+ staining but a persistence of some GFAP+ cells were classified as severe glaucoma. These data suggest a phasic stratification with disease where maximal glial activation is seen in the moderate stage.

### Changes in immunohistochemical markers stratified by glaucoma severity

After implementation of clustering and definition of the three distinct glaucoma groups, we reassessed our data based on this refined system. We observed a progressive loss in RBPMS+ RGCs and NEFL staining in the temporal retinas with increasing glaucoma severity (**Fig. 9A, B**). GFAP+ cell density was severity-dependent, demonstrating maximal glial activation with moderate severity (**Fig. 9C**). On the other hand, IBA1+ immunoreactivity increased in the moderate and severe glaucoma groups with large variability but did not drastically decrease in severe glaucoma (**Fig. 9D**).

**Figure 9:**
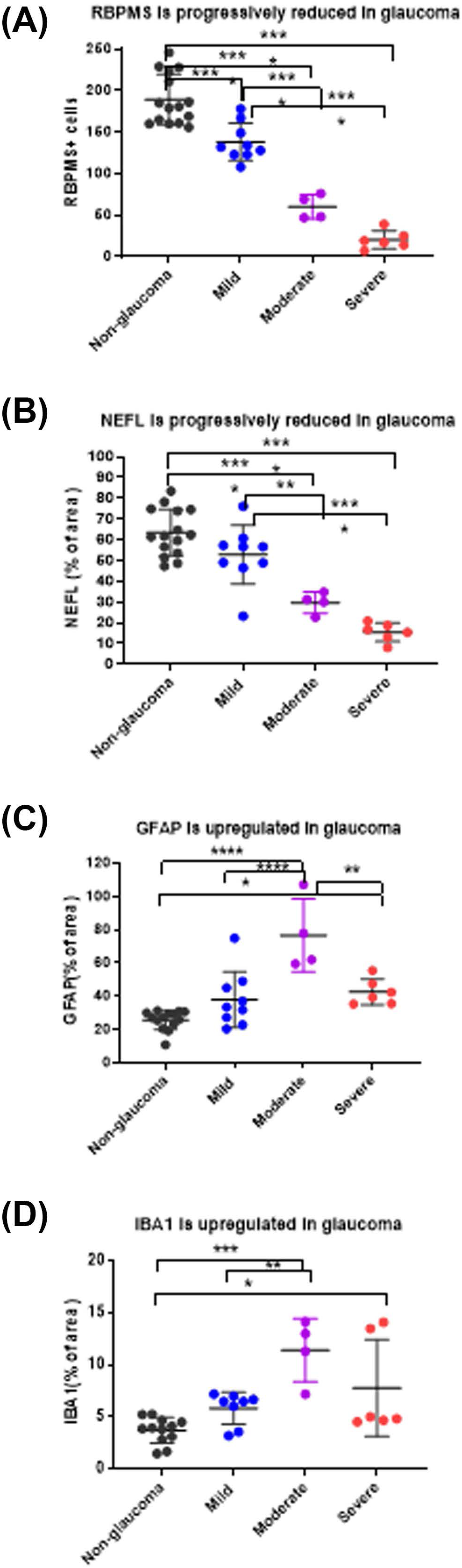
Post-hoc analysis after stratification of glaucomatous donors. Progressive RGC degeneration and increased glial reactivity were observed in the retina with disease severity. **(A)** Loss of RBPMS+ RGC cells, **(B)** loss of axons (NEFL), **(C)** increased astrocytic activation (GFAP), and **(D)** microglia/macrophage activation (IBA1) was stratified by disease state. Horizontal bars indicate mean ± standard deviation. *p<0.05, **p<0.01, ***p<0.001, ****p<0.0001 One-way Analysis of variance (ANOVA) followed by Tukey’s multiple comparison test.

## DISCUSSION

Clinically, glaucoma severity is subjectively graded using structure-function analysis, predominantly by compiling a composite of fundus images, vertical cup-to-disc ratios (VCDR), OCT images, and visual field metrics. To the best of our knowledge, there are no unified standards used to define disease severity, although significant efforts are underway to address this [1]. While in-life longitudinal assessment of glaucoma severity is ideal and can monitor a patient’s disease progression, it is challenging to correlate progression with specific molecular changes at the sites of pathology in the neural retina and optic nerve in the same patient, owing to the inability to perform successive biopsies whilst maintaining vision and function. Thus, researchers have increasingly turned to using post-mortem eyes obtained from eye banks to identify biomarkers and new therapeutic targets. Here, the challenges are different, in that longitudinal clinical imaging and detailed medical histories are typically not provided and rely heavily on self-disclosure of ocular disease history by the donor’s family. Conversely, glaucomatous changes such as retinal thinning and severe optic nerve cupping can be found post-mortem in eyes with undiagnosed glaucoma. Thus, there is a need to better stratify donor samples with incomplete histories to better understand the utility of the tissues for mechanistic investigation. In this study, we used multiparametric histological assessments, focusing on molecular markers, to classify glaucoma severity in post-mortem donor tissues.

There is currently no consensus for histology-based morphometric, RNA, and protein markers that are accurate and predictive diagnostics for glaucoma stage. In addition, death-to-preservation time and fixation procedures can create confounding artifacts. Thus, relying only on measurements of physical features could lead to a less accurate and comprehensive assessment of disease stage. We selected a combination of immunohistologic measurements in the neural retina and ONH that we reasoned would give key estimates of severity: (1) the number of RBPMS+ RGCs, to assess remaining RGC soma; (2) neurofilament staining as measured by NEFL, to assess the extent of axonal and dendritic loss; (3) the number of IBA1+ cells, to assess microglia/macrophages and estimate the degree of innate inflammation in the tissue, and; (4) the number of GFAP+ cells, to assess astrocytes and estimate the degree of gliosis in the tissue. Individual assessment of each marker showed a decrease of RBPMS and NEFL staining and an increase in IBA1 and GFAP staining in glaucoma samples, as expected. A factor to consider when grading is the consistency in available tissue area. While the retinal regions could be consistently normalized, we observed significant variability in cupping and the amount of optic nerve and ONH tissue between glaucomatous donor sections, which could impact normalization between donors. To minimize the effects of this potential confounder, we focused our quantitative assessments using retinal measurements. Combining all four quantitative measurements in a multiparametric approach, we were able to stratify our glaucoma donor samples into three distinct groups of mild, moderate, and severe. This type of stratification was not attainable using donor records or individual markers. Our conceptual grading approach is outlined in **Fig. 10**.

**Figure 10:**
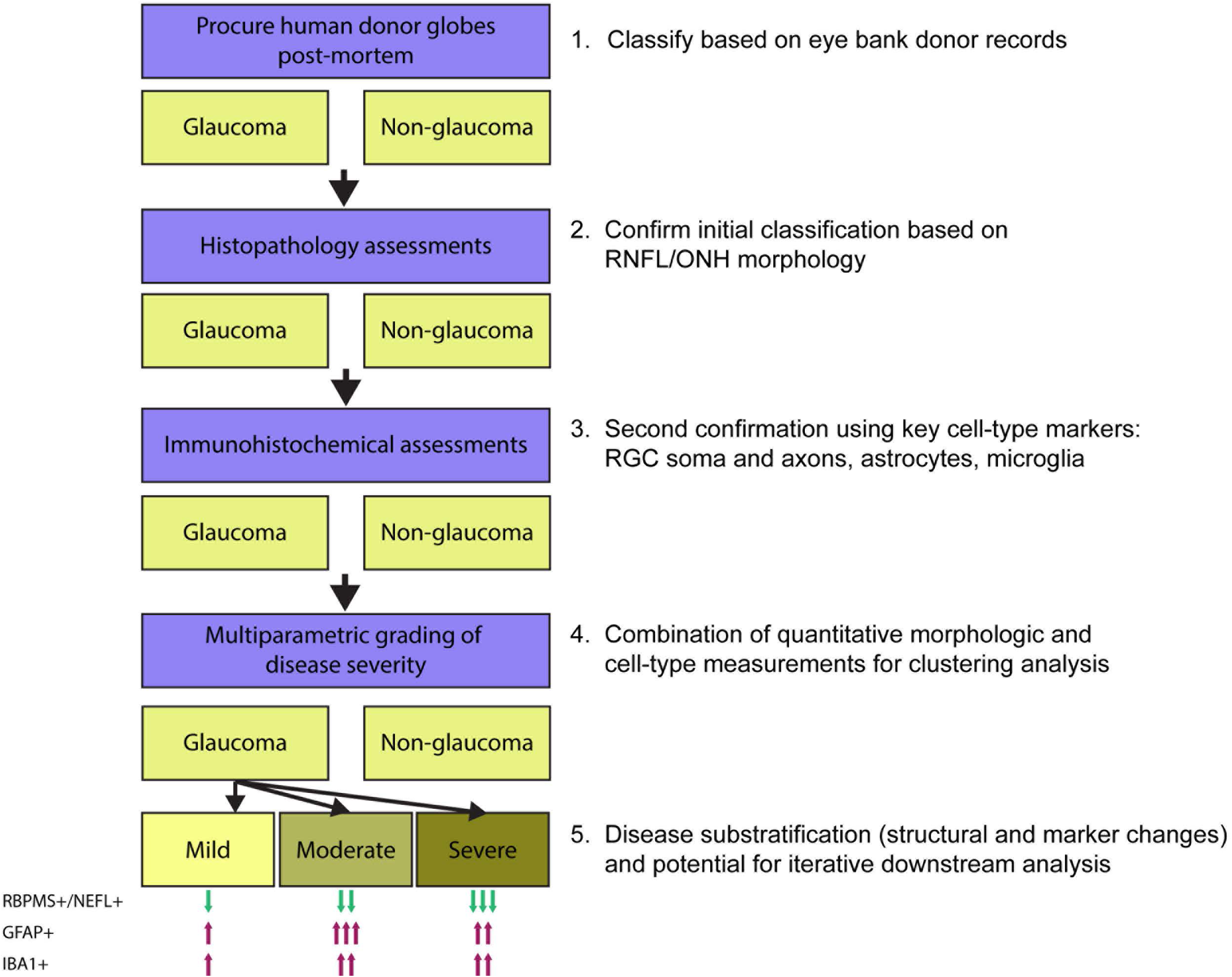
Schematic of donor eye grading workflow. Conceptual outline of steps in acquisition and grading of post-mortem donor eyes, accompanied by qualitative descriptions of protein marker stain changes from non-glaucoma samples in each severity grade. Note that the evaluation of protein markers is confined to the temporal retina. One arrow indicates a mild/minimal change, two arrows indicate a moderate change, and three arrows indicate a severe/maximal change. The predominant/distinguishing features of each grade are: mild – pre-mortem history of glaucoma; moderate – maximal GFAP positivity in temporal neural retina; severe – maximal loss of RBPMS and NEFL positivity in temporal retina.

Glia (microglia, astrocytes and Müller glia) play roles either in restoring homeostasis or exacerbating disease conditions in neuropathies. Notable microglial changes in neuroinflammatory conditions include an increase in their numbers, IBA1+ expression, and upregulation of complement factors such as C1Q and C3. Microglial changes are time dependent in relation to their effects on RGC-astrocyte interactions and RGC survival. Early microglial changes likely involve synaptic pruning via complement deposition, followed by more persistent pro-inflammatory phenotypes that can indirectly promote neuroinflammation through engaging reactive astrocytes to secrete cytokines and vasoactive peptides such as endothelin-1 [9, 10, 28]. We observed a 100% increase in IBA1+ cells in glaucomatous donor retinas when compared to non-glaucomatous donors, with enrichment in the RGC and inner plexiform layers. The increase in IBA1+ cells was most pronounced in the moderate and severe glaucoma groups. In contrast, while the number of GFAP+ cells also increased in glaucoma, there was not a statistically significant correlation between GFAP+ and IBA+ expression across glaucoma samples after stratification into stages, suggesting a temporal disconnect between microglial and astrocyte activation, proliferation, and modulation over the course of disease progression. Alternatively, astrocyte activation could be different in the ONH, given that predominant glial activation changes in glaucoma have been observed in the unmyelinated region of the optic nerve. We did not evaluate molecular changes in the ONH in this study, as its size varied between donors, and our initial classification of disease was based on the presence or absence of glaucoma from donor medical records. The implication of the temporal disconnect between activation of different glial cells is as yet unclear, and further mechanistic studies may be required to delineate their specific roles and spatial distribution (if any) in initiation and progression of neurodegeneration. A recent literature meta-analysis by Salkar et al [29] of molecular changes in human glaucomatous post-mortem retina and ON tissues confirmed associations of glial cell activation and neuroinflammation. This approach yielded a comprehensive list of neuroinflammatory participants in astrocyte activation and the possibility of interplay with other glial cells (microglia and Müller glia). The authors used a standardized mean differences quantitation to identify key players that favored neuroinflammation in glaucoma, including GFAP, p-ERK, nitrotyrosine, endothelin B receptor, and RhoA [29].

The explosion of highly multiplexed transcriptomics and proteomics platforms is enabling greater resolution of molecular markers in human tissues. For example, a recent study [30] identified 37 clusters and subclusters of distinct cell types in the optic nerve, optic nerve head, retinal pigmented epithelium and peripapillary sclera of non-glaucomatous donor globes. These findings offer insights into expression profiles that define cell subtypes, and computational approaches can be applied to these datasets to infer cell-cell interactions. The potential to overlay these detailed molecular phenotypes on top of histopathological grading of glaucoma severity will likely be important in identifying disease and cell-type specific targets that are associated with distinct disease stages. As an example, it has been noted that the cell-type specific expression patterns of glaucoma risk genes (including primary open-angle and normotensive glaucoma) and genes that confer susceptibility to increased IOP but not glaucoma are distinct and different. Specifically, IOP susceptibility genes are reported to be expressed in vascular cells such as Schlemm’s canal endothelia, whereas POAG risk genes are primarily expressed in the RPE, retinal and ON astrocytes, Muller glia, and oligodendrocytes. While all of these studies were performed using non-glaucomatous donor tissues, one could consider conducting a similar assessment, accompanied by multiplexed spatial transcript and protein profiling, on samples graded with the severity index from our current study. This approach could identify novel targets and mechanisms of action and lead to new therapeutic hypotheses. For example, impairment in microglial autophagy has been noted in neurodegeneration and aging [31]. Whether a similar impairment exists in glaucoma is unclear but could be assessed via a combination of spatial transcriptomics and immunohistochemistry along with anatomical assessment of disease severity. The differential expression profiles of microglial subtypes are indicative of putative pro- and anti-neuroinflammatory roles, particularly when considering surface receptor profile changes. For instance, the CX3CR1 receptor appears to play a role in eliminating dead or dying cells like RGCs, and antagonism of this receptor exacerbates RGC loss via microglial activation in preclinical glaucoma models [32, 33]. Microglial adenosine A2A receptor or A3 receptor antagonism has been shown to be neuroprotective in glaucomatous rodent *in vitro* and *in vivo* models [34, 35]. Therefore, these examples suggest that profiling of cell surface receptor mRNA transcripts and protein levels may yield further insights into spatial and disease-specific stratification when combined with multiparametric indexing of glaucomatous donor globes.

There are several gaps and limitations in our approach that could be addressed by further experimentation and validation. First, we currently lack a sufficient number of samples that could make up an independent validation cohort, and have not yet performed a confirmatory exercise. Independent replication of our grading approach by other groups would be an important test of whether the results from donor samples obtained from one site are robust and reproducible. Second, as previously mentioned, there is no current consensus on specific individual or combinations of protein markers, or the precise magnitude of change in these markers, that can be used to reliably classify glaucoma disease severity. Third, the complexity of gliosis and neuroinflammation suggests that reliance on a few RNA or protein markers may not be sufficient to thoroughly describe the complexity of cell phenotypes in disease, especially without spatial resolution. Identification of additional markers through various technologies and subsequent incorporation into a grading scheme could further refine our system, or potentially alter the classification scheme. Finally, the statistical approaches used to sub-stratify our samples could be further refined, or additional methods incorporated. We also note that some of the statistical methods used may not be immediately accessible to some researchers and pathologists. Our conceptual grading framework provides a starting point for discussion and will enable further experimentation and refinement, with the ultimate goal of building a consensus methodology for histology-based grading of glaucoma samples. As consensus is refined, development of automated classification systems could be pursued.

Looking ahead, there is still a disconnect between the types of measurements that can be made in post-mortem ocular samples and from live patients. It is challenging to obtain donor globes from large-scale healthcare data curation efforts with a rich set of longitudinal imaging and functional measurements. In the rare cases where this is possible, molecular assessments will undoubtedly be terminal. To correlate molecular and clinically observable changes across the course of glaucoma progression, bridging studies and efforts will need to be made to link measurements such as structure (VCDR, in-life RNFL thickness), function (e.g. HVF), genetics, and disease-associated molecular changes at cellular resolution. With recent developments in AI algorithms and available training datasets, it is more feasible that post-mortem histological assessments could be correlated with similar in-life measurements. For example, rapid grading of a larger number of clinical images of optic nerve heads can enable prediction of progression [36]. Streamlining donor sample acquisition with detailed clinical information regarding glaucoma diagnosis and progression will enable higher resolution, high-throughput multi-omic approaches to understand mechanisms of disease progression. If a subset of these types of images can be linked to post-mortem assessments from the same patients, followed by comprehensive RNA and protein profiling, these efforts could serve as the basis for a curated dataset for AI-based scoring of donor globes with or without longitudinal data. Such a resource could greatly accelerate our ability to understand glaucoma progression at cellular and molecular resolution and develop the next wave of neuroprotective therapies.

## ACKNOWLEDGEMENTS

We thank David Ammar, Nicholas Sprehe, Jason Woody and the team at Lion’s Eye Institute for Transplant and Research (now Lion’s World Vision Institute) for their partnership in donor tissue procurement, and Chia-Ling Huang, Chenying Guo and Nathaniel Kirkpatrick for advice and support. Lastly, we would like to thank the donors and their families for their generous gift to research

---

**Supplementary Figure 1: Schematic representation of regions utilized for quantitative histological analysis.** Retinal thickness measured on H&E slides. Thickness of the Ganglion Cell Complex (GCC) including the retinal nerve fiber layer (RNFL) and retinal ganglion cell (RGC) layer plus inner plexiform layer was measured. 30-40 measurements were conducted in the 2 mm range of temporal peripheral retina (beginning from the single layer from the macula). Cell quantification: RBPMS+ cells in the 5 mm range of temporal retina were counted. Quantification of NEFL by area: The percentage area of the positive stained signal (%) of the total area measured in 2 mm range was calculated, vessels were excluded.

**Supplementary Figure 2: Variability observed between donors with a medical history indicating glaucoma. (A)** The RGC layer was lost in both temporal and nasal macular regions in sections from this donor. **(B)** A second, different donor, exhibiting localized loss of RGCs and a reduction in RGC density in the central and temporal macula, respectively.

**Supplementary Figure 3: Representative images of co-staining of IBA1+ and RBPMS+ cells in glaucomatous donors.** Peripheral retinas immunolabeled with RGC (RBPMS; red arrow) and microglial (IBA1; blue arrow) cell markers demonstrated that both cell types exhibited a close (proximity) association with each other (n=3).

**Supplementary Figure 4:** Receiver operating characteristic curve analysis was performed to distinguish between false positive or false negative differences between values for glaucomatous and non-glaucomatous donors. ROC AUCs showed that expression levels of (A) RBPMS, (B) NEFL and (C) GFAP demonstrate significant differences between glaucomatous and non-glaucomatous donor tissues with RBPMS serving as the best indicator for disease discrimination.

