## Supplementary material for "Multiparametric grading of glaucoma severity by histopathology can enable post-mortem substratification of disease state": Table 1

| **Donor #** | **Disease state** | **Age** | **Sex** | **Race/Ethnicity** | **Cause of death** |
| --- | --- | --- | --- | --- | --- |
| 1 | Non-glaucoma | 94 | male | Caucasian | COPD |
| 2 | Non-glaucoma | 92 | male | Caucasian | Cardiac arrest. |
| 3 | Non-glaucoma | 84 | female | Caucasian | Multiple Myeloma |
| 4 | Non-glaucoma | 84 | male | Caucasian | Acute Cardiac Crisis |
| 5 | Non-glaucoma | 77 | male | Caucasian | Sepsis |
| 6 | Non-glaucoma | 84 | male | Caucasian | Severe PVD |
| 7 | Non-glaucoma | 73 | male | Caucasian | Leukemia |
| 8 | Non-glaucoma | 78 | male | Caucasian | CHF, CVA |
| 9 | Non-glaucoma | 86 | male | Caucasian | Pneumonia |
| 10 | Non-glaucoma | 81 | male | Caucasian | Prostate cancer, Diabetes |
| 11 | Non-glaucoma | 84 | male | Caucasian | End stage lung cancer |
| 12 | Non-glaucoma | 83 | female | Caucasian | Lung cancer with mets, Pneumonia |
| 13 | Non-glaucoma | 83 | female | Caucasian | Trauma, Hospice. |
| 14 | Non-glaucoma | 68 | male | Caucasian | Cellulitis of Leg, Severe PVD, Acute Severe Anemia |
| 15 | Non-glaucoma | 93 | female | Caucasian | Pneumonia |
| 16 | Non-glaucoma | 79 | female | Caucasian | Anemia, HTN, COPD, |
| 17 | Glaucoma | 73 | male | Caucasian | prostate CA |
| 18 | Glaucoma | 86 | male | Caucasian | Metabolic Encephalopathy |
| 19 | Glaucoma | 87 | male | Caucasian | Skin Cancer |
| 20 | Glaucoma | 97 | male | Caucasian | Acute cardiac crisis |
| 21 | Glaucoma | 76 | female | Caucasian | MRSA |
| 22 | Glaucoma | 65 | female | Caucasian | COPD, Pneumonia |
| 23 | Glaucoma | 79 | female | Caucasian | Pneumonia |
| 24 | Glaucoma | 92 | female | Caucasian | Acute myeloid leukemia |
| 25 | Glaucoma | 78 | female | Caucasian | Pneumonia |
| 26 | Glaucoma | 77 | male | Caucasian | COPD, Pneumonia |
| 27 | Glaucoma | 93 | male | Caucasian | Pancreatic Cancer |
| 28 | Glaucoma | 88 | male | Caucasian | Renal failure |
| 29 | Glaucoma | 80 | female | Caucasian | Acute myeloid leukemia; |
| 30 | Glaucoma | 68 | female | Black | Ichthyophthirius multifiliis |
| 31 | Glaucoma | 83 | male | Asian | Fecal impaction, Weakness, Pleural effusion |
| 32 | Glaucoma | 97 | female | Caucasian | COPD, respiratory failure |
| 33 | Glaucoma | 90 | male | Caucasian | Acute cardiac event |
| 34 | Glaucoma | 85 | female | Caucasian | Acute kidney injury |
| 35 | Glaucoma | 77 | female | Caucasian | COPD, respiratory failure |
| 36 | Glaucoma | 71 | male | Caucasian | COPD, Pneumonia |
| 37 | Glaucoma | 70 | male | Caucasian | COD, Dementia |
| 38 | Glaucoma | 78 | female | Caucasian | intracerebral bleed |
