## Supplementary material for "Multiparametric grading of glaucoma severity by histopathology can enable post-mortem substratification of disease state": Table 2

Table 1: IHC antibody information

| Antibody | Vendor | Cat# | Host | Stock concentration (µg/ml) | Dilution factor |
| --- | --- | --- | --- | --- | --- |
| RBPMS | Novus | NBP2-20112 | Rabbit polyclonal | 1000 | 1:800 |
| NFL | Proteintech | 12998-1-AP | Rabbit polyclonal | 600 | 1:1000 |
| GFAP | Abcam | Ab7260 | Rabbit polyclonal | 1000 | 1:6000 |
| IBA1 | Wako | 019-19741 | Rabbit polyclonal | 1000 | 1:5000 |
