## Supplementary figures and images for "Multiparametric grading of glaucoma severity by histopathology can enable post-mortem substratification of disease state"

### Supplemental Figure 1

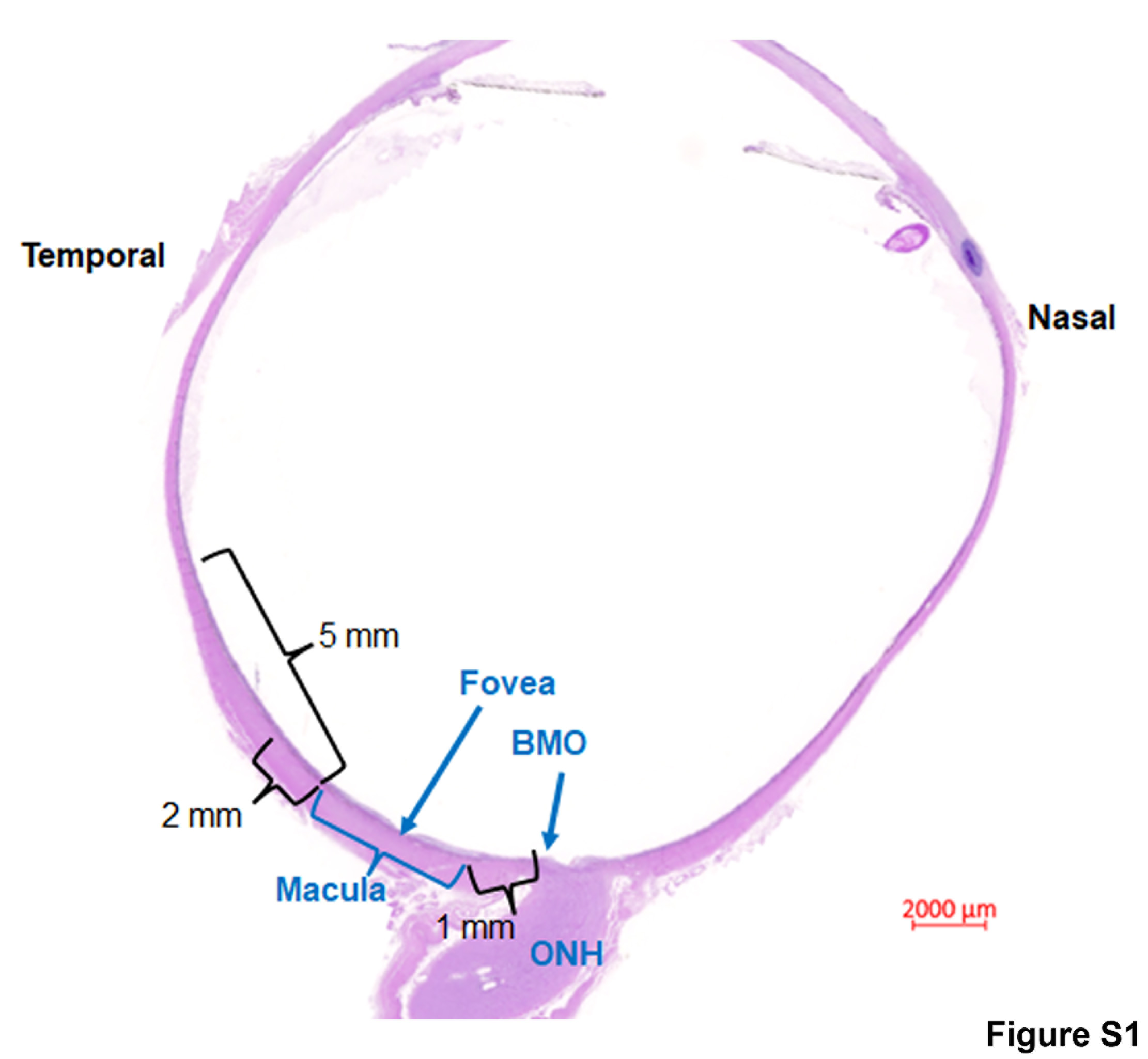

### Supplemental Figure 2

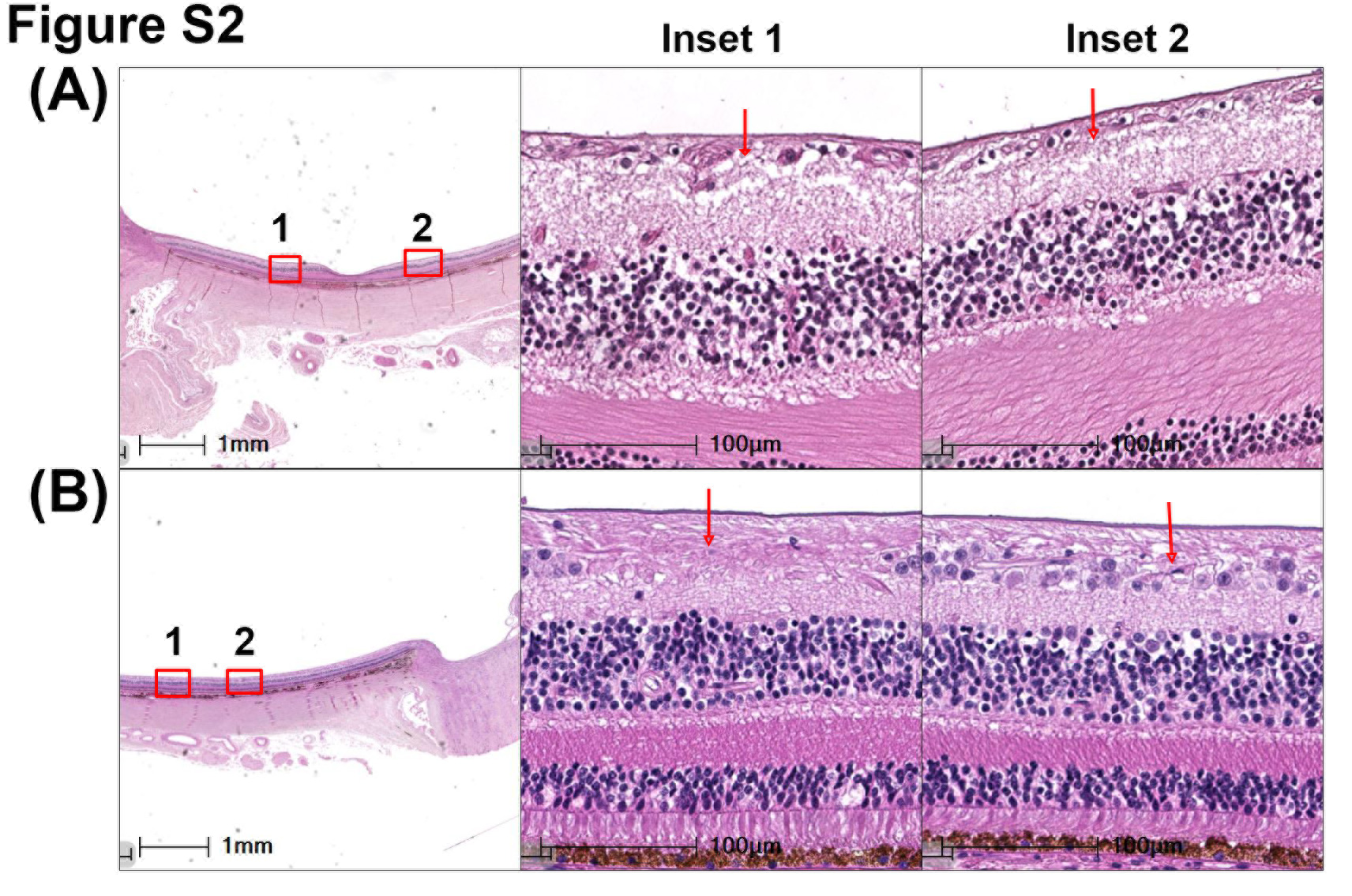

### Supplemental Figure 3

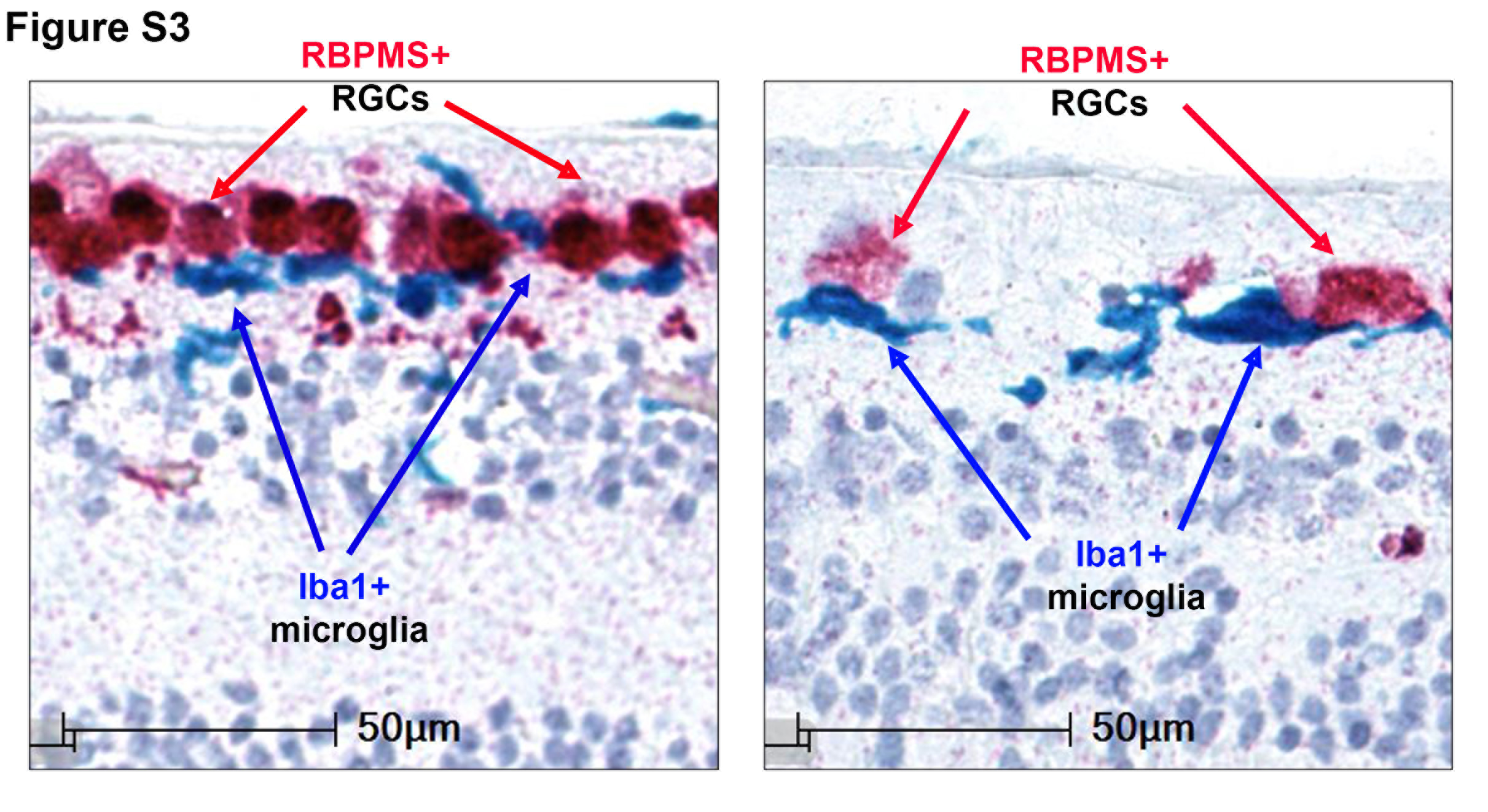

### Supplemental Figure 4

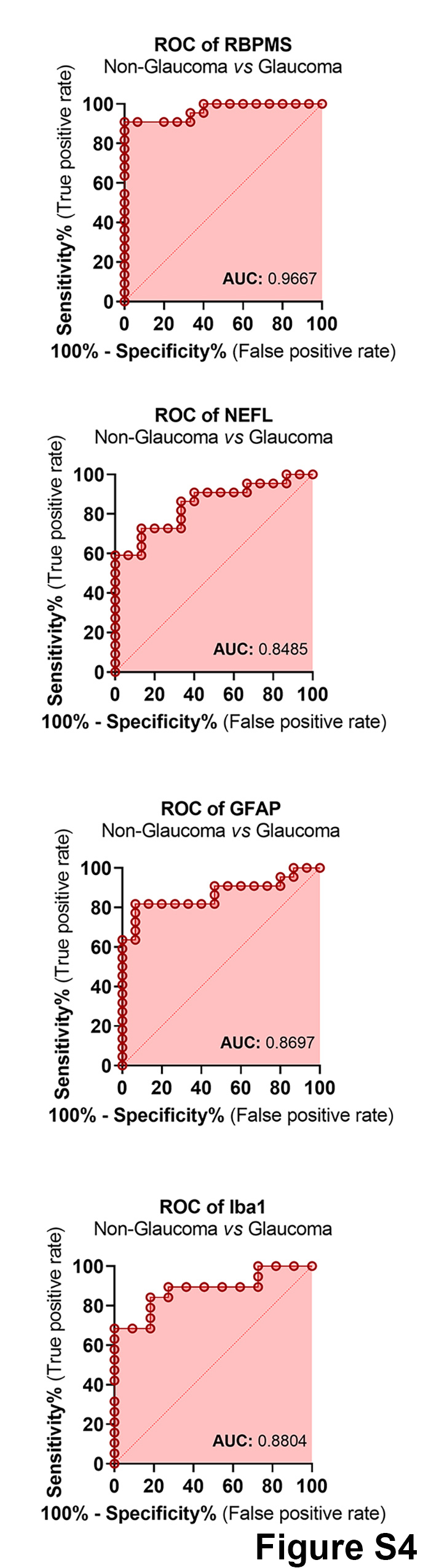
